# The scaffolding protein Flot2 regulates cytoneme-based transport of Wnt3 in gastric cancer

**DOI:** 10.1101/2022.01.07.475396

**Authors:** Daniel Routledge, Sally Rogers, Hassan Ashktorab, Toby J Phesse, Steffen Scholpp

**Affiliations:** Living Systems Institute, School of Biosciences, College of Life and Environmental Sciences, University of Exeter, Exeter, UK; Department of Medicine, Howard University, 2041 Georgia Ave, Washington, DC 20060, US; European Cancer Stem Cell Research Institute, Cardiff University, Cardiff, UK

## Abstract

The Wnt/β-catenin signalling pathway regulates multiple cellular processes during development and many diseases, including cell proliferation, migration, and differentiation. Despite their hydrophobic nature, Wnt proteins exert their function over long distances to induce paracrine signalling. Recent studies have identified several factors involved in Wnt secretion, however, our understanding of how Wnt ligands are transported between cells to interact with their cognate receptors is still debated. Here, we demonstrate that gastric cancer cells utilise cytonemes to transport Wnt3 intercellularly to promote proliferation. Furthermore, we identify the membrane-bound scaffolding protein Flotillin-2 (Flot2), frequently overexpressed in gastric cancer, as a regulator of these cytonemes. Together with the Wnt co-receptor and cytoneme initiator Ror2, Flot2 determines the number and length of Wnt3 cytonemes in gastric cancer. Finally, we show that Flot2 is necessary for Wnt8a cytonemes during zebrafish embryogenesis, suggesting a conserved mechanism for Flot2-mediated Wnt transport on cytonemes in development and disease.

## Introduction

Wnt/β-catenin signalling activity has been well-characterised in the glands of the gastrointestinal epithelium (Clevers, 2013). In these intestinal crypts, the expression of Wnt/β-catenin target genes occurs in a gradient, with the highest activity at the base of the crypt (Muñoz *et al*., 2012). Here, Paneth cells are located between the intestinal stem cells and regulate proliferation by producing high levels of Wnt3 (Farin *et al*., 2012; Sato *et al*., 2011). Genomic analysis suggests that in about 50% of gastric tumours, the Wnt/β-catenin pathway is deregulated (Koushyar *et al*., 2020).

In many gastric carcinomas, Wnt3 expression is elevated compared with normal gastric epithelium, leading to increased proliferation in the tumour mass (Voloshanenko *et al*., 2013; Wang *et al*., 2016). In development, Wnt proteins can spread over a distance of many cells despite their hydrophobic nature (Routledge and Scholpp, 2019). However, it is unclear how lipid modified Wnt3 is disseminated in gastric tumour tissues.

Recently, thin and actin-rich membranous protrusions known as cytonemes have been identified as crucial for transporting essential signalling components between cells (Kornberg and Roy, 2014; Zhang and Scholpp, 2019). For example, in zebrafish embryogenesis, cytonemes are highly dynamic and have been reported to transport Wnt components (Stanganello *et al*., 2015), whilst in the chick limb bud cytonemes transport Shh (Sanders *et al.,* 2013). Wnt/PCP signalling in the source cell facilitates the emergence of long and stable Wnt-transporting cytonemes, promoting paracrine Wnt signal activation (Mattes *et al*., 2018; Brunt *et al*., 2021).

Similarly to Wnt3, Flot2 is linked to tumour development and proliferation, and its expression is increased in gastric cancer (GC) (Cao *et al*., 2013; Zhu *et al*., 2013). Flotillins are highly conserved, membrane-bound scaffolding proteins, which cluster in detergent-resistant, cholesterol-rich regions of the membrane (Lang *et al*., 1998; Stuermer *et al*., 2001). These Flotillin microdomains act as platforms to facilitate protein-protein interactions and regulate growth factor signalling (Banning *et al*., 2014). In *Drosophila,* Flotillin-2/Reggie-1 (Flot2) promotes long-range spreading and signalling of the lipophilic ligands, Wg and Hh (Katanaev *et al*., 2008) and Flot2 can promote the formation of Hh cytonemes (Bischoff et al., 2013). In cancer cell lines, Flot2 also facilitates the formation of filopodia-like protrusions via direct regulation of the actin cytoskeleton (Neumann-Giesen *et al*., 2004; Gauthier-Rouvière *et al*., 2020).

In this study, we found that human GC cells use cytonemes to disseminate Wnt3 proteins intercellularly. We further detected that these Wnt3-bearing cytonemes are dependent on Flot2 function. Our data suggest that Flot2 function is required for intracellular trafficking and signalling of the Wnt co-receptor Ror2, a key regulator of cytoneme formation. In accordance with our results in GC cells, we identified Flot2 as a regulator of Wnt8a cytoneme emergence in zebrafish development, suggesting a conserved role for Flot2 in Wnt cytoneme emergence in vertebrates.

## Results

### Gastric cancer cells utilise cytonemes for paracrine Wnt3 signalling

First, we examined the gastric adenocarcinoma cell lines MKN-28, MKN-7, and AGS and compared these to a gastric epithelial cell line HFE-145 for their potential to form actin-based protrusion. All GC cells form many dynamic actin-based filopodia, and AGS cells display the longest filopodia (Fig. 1a, b). Next, we asked if GC cells use these filopodial protrusions to mobilise Wnt3 ligands. Using our advanced filopodia fixation and immunohistochemistry (IHC) protocol (Rogers and Scholpp, 2021), we observed that HFE-145 have numerous protrusions, which are decorated with Wnt3 (Fig. 1c). In the cancer cell lines, Wnt3 localisation was enriched on these filopodia and more often detectable (Fig. 1c, d). Specifically, the AGS line shows an increased number of Wnt3 cytonemes (Fig. 1d) and we, therefore, used AGS in the further analysis. Over-expression of Wnt3 in AGS led to an even further increased Wnt3 staining on more filopodia (Fig. 1d, e). After treatment of AGS cells with the porcupine (PORCN) inhibitor IWP2, we could not detect Wnt3 localisation at the plasma membrane and on filopodia (Fig. 1f), suggesting that PORCN-dependent palmitoylation is essential for localisation of Wnt3 on these actin-based protrusion. Similar to the endogenous protein, we found that Wnt3-mCherry is transported along filopodia and localises to their tips (Fig. 1g, Supp. Video 1). We also assessed the localisation of Myosin-X (MyoX), a filopodial motor protein involved in Shh cytoneme function by IHC and observed strong localisation on filopodia tips (Fig. 1h) (Bohil *et al*., 2006; Hall *et al*., 2021). Similarly, an IHC analysis showed that the Wnt transporter Evi/Wntless localises to these filopodia, suggesting a possible role in transporting Wnts along these protrusions (Fig. 1i). We conclude that Wnt3 can be loaded on filopodia-like protrusion, which we termed cytonemes, based on their definition as signalling filopodia (Kornberg 2014; Mattes and Scholpp, 2018).

**Figure 1.**
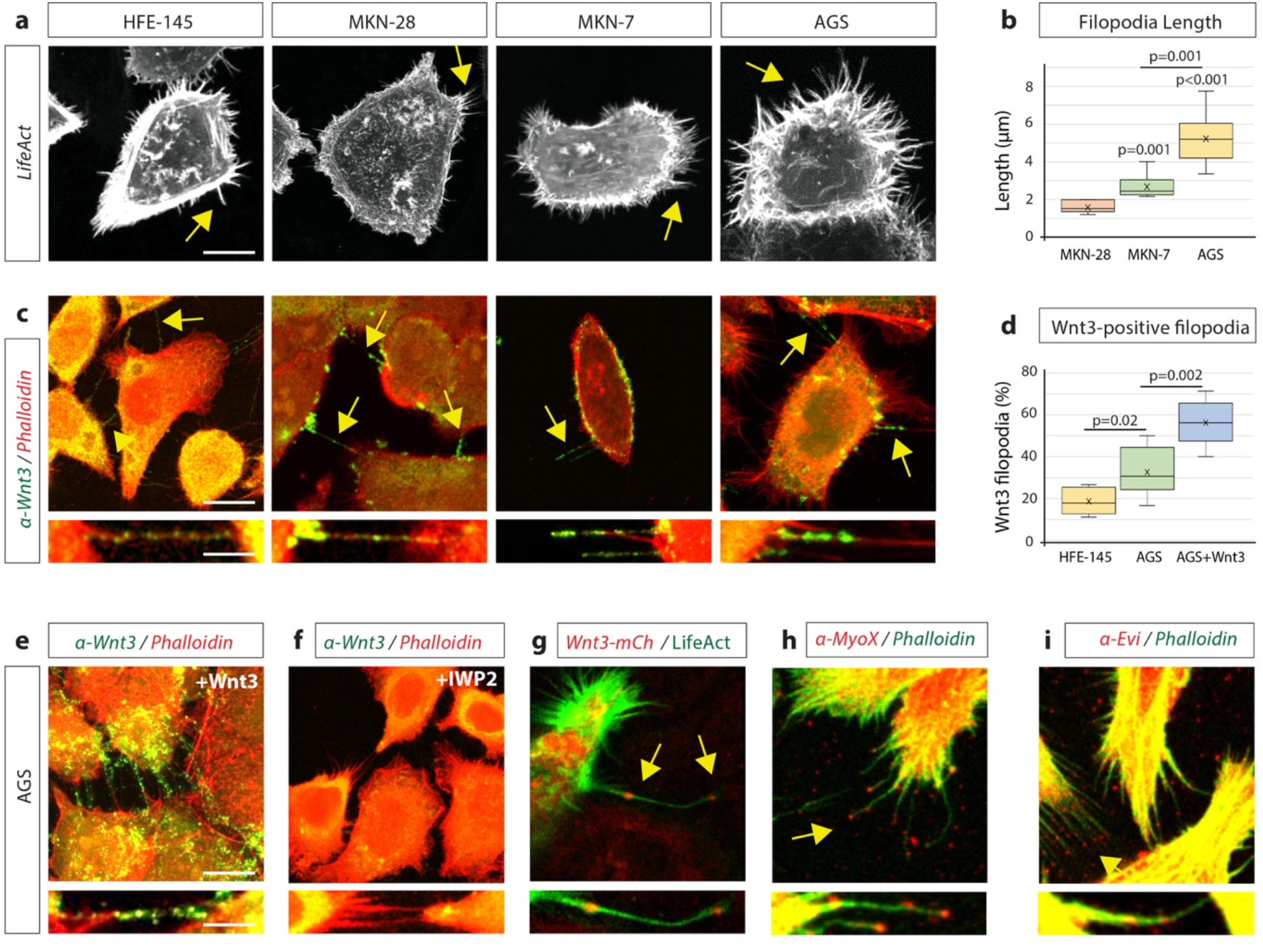
Gastric epithelial normal and cancer cell lines utilise cytonemes to transport Wnt3 intercellularly. **a,** Confocal images of normal gastric epithelial cell line (HFE-145) and gastric cancer cell lines (MKN28, MKN7, and AGS) expressing LifeAct-GFP to visualise actin-based structures. Examples of filopodia are indicated by yellow arrows. **b**, Quantification of filopodia length in gastric cancer cell lines MKN28, MKN7 and AGS (n = 7, 8, 25; n = number of cells). Significance calculated by Student’s t-test. **c,** Immunofluorescent (IHC) images of HFE, MKN28, MKN7, and AGS, stained with antibodies against endogenous Wnt3 (green) and actin (iFluor594, red). Scale bar 10µm. High-magnification images indicate an example of a Wnt3-bearing cytonemes. Scale bar 2.5µm. **d,** Quantification of Wnt3-positive filopodia in gastric epithelial (HFE-145) and cancer (AGS) cells as a percentage of total filopodia (n = 6, 8, 6; n = number of cells). Significance calculated by Student’s t-test. **e,** IHC images of AGS cells overexpressing Wnt3 and antibody stained for Wnt3 (green) and actin (iFluor594, red). Scale bar 10µm. High-magnification images highlight cytonemes. Scale bar 2.5µm. **f,** IHC images of AGS cells treated with the Porcupine inhibitor IWP2 (100µM, 48 hrs) and stained with an antibody against Wnt3 (green) and actin (iFluor594, red). **g,** Live confocal cell imaging of AGS cells expressing Wnt3-mCherry and LifeAct-GFP. Cytoneme-localised Wnt3-mCherry highlighted by yellow arrows. **h-i,** IHC images of AGS cells stained with antibody stained for (**h**) Myosin-X (MyoX) and (**i**) Evi/Wntless (red). FITC phalloidin labels actin (green).

### Wnt3 cytonemes regulate paracrine Wnt/β-catenin signalling in gastric cancer cells

Next, we assessed the functional impact of Wnt3 cytonemes on paracrine Wnt signalling in gastric cancer cells by performing a co-cultivation assay of Wnt-producing AGS cells and Wnt-receiving HFE-145 cells (Fig. 2a). To specifically block cytoneme formation, we used a dominant-negative mutant of the Insulin Receptor tyrosine kinase Substrate p53 (IRSp53). IRSp53 is a multidomain BAR protein, which binds active Cdc42 and N-Wasp to promote filopodia formation (Kast *et al*., 2014). Here, the mutated protein IRSp53^4K^ blocks filopodia formation (Supp. Fig. 1a,b) (Meyen *et al*., 2015) without interfering with autocrine Wnt signalling (Stanganello *et al*., 2015; Brunt *et al*., 2021). IRSp53^4K^-GFP significantly reduced paracrine Wnt SuperTOPFlash (STF) reporter activation (Fig. 2b, c). Transfection of Wnt3 in the source cells doubled STF reporter activity in the co-cultivated HFE-145 cells, which could be attenuated significantly by the co-transfection of IRSp53^4K^-GFP. Our data suggest that cytonemes are vital for paracrine Wnt/β-catenin signal activation. Furthermore, we conclude that the majority of Wnt3 protein is disseminated via IRSp53-dependent cytonemes in gastric cancer cells.

**Figure 2.**
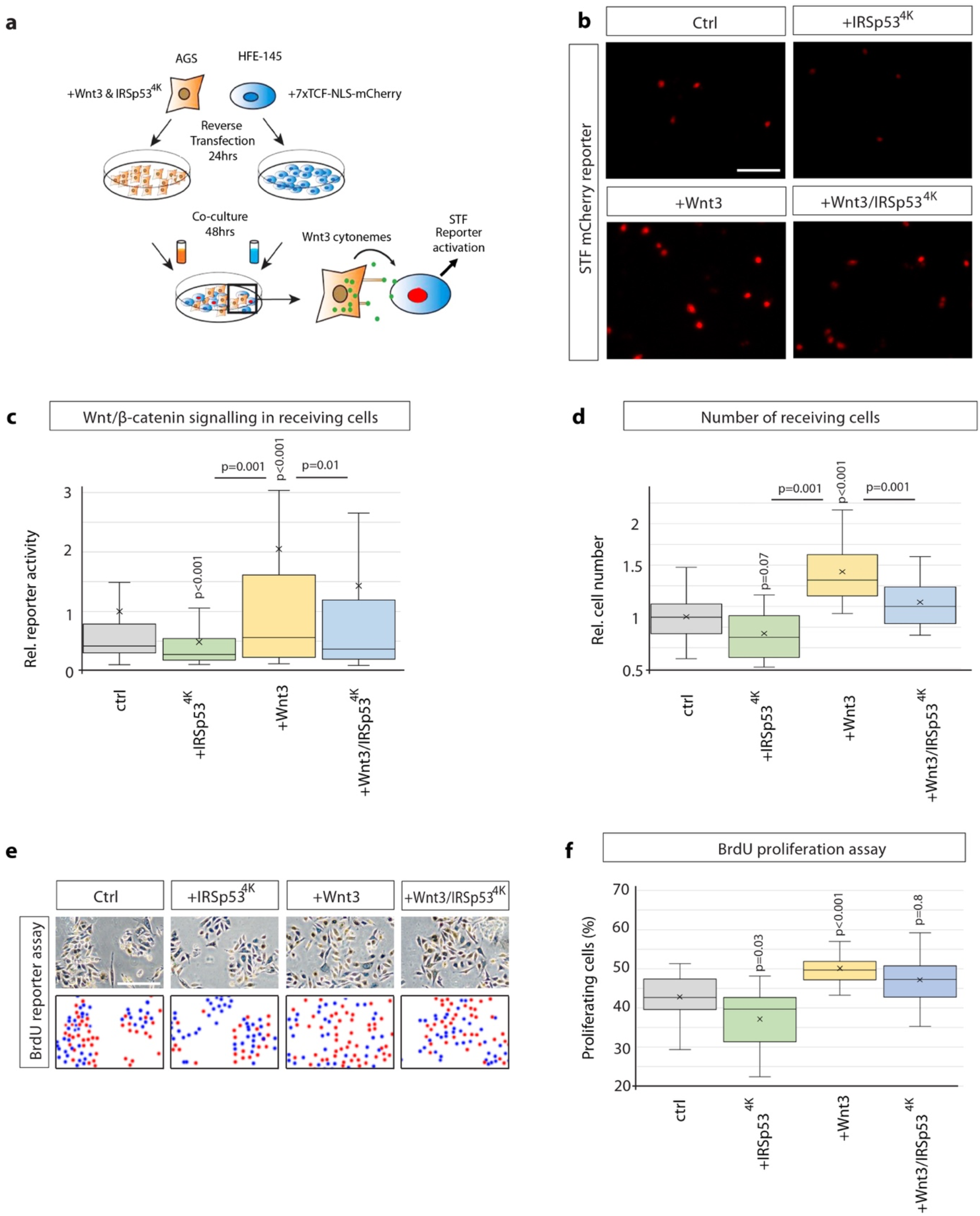
Wnt3 cytonemes regulate paracrine Wnt/β-catenin signalling and proliferation. **a,** Experimental protocol for measuring paracrine Wnt signalling activation. HFE cells expressing the SuperTOPFlash (STF) reporter, 7xTCF-NLS-mCherry, were co-cultivated with AGS cells expressing indicated constructs. Fluorescence of STF mCherry reporter was measured after 48 hrs and compared to untransfected control. **b,** Representative images of STF reporter fluorescence for indicated conditions. Scale bar 100µm. **c,** Quantification of STF mCherry reporter fluorescence in HFE cells co-cultivated with AGS (n per condition = 322, 394, 258, 275). **d,** Relative number of HFE cells per image after co-cultivation with AGS cells expressing indicated constructs. Significance calculated by Student’s t-test with Bonferroni correction for multiple comparisons. (n per condition = 28, 26, 27, 17; n = number of images). **e,** Representative images of proliferating, BrdU-stained (red); co-cultivated AGS and HFE-145 cells, as described in (a). Cells were counterstained with haematoxylin (blue dots). Scale bar 100µm. **f,** Quantification of BrdU-stained cells as a percentage of the population. Significance calculated by student’s t-test with Bonferroni correction for multiple comparisons. (n per condition = 20, 20, 20, 20; n = number of images).

The Wnt/β-catenin signalling pathway is essential for proliferation and self-renewal of both cancer stem and progenitor cells (Koushyar *et al*., 2020). To assess the impact of paracrine Wnt signalling on cell proliferation, the number of HFE-145 cells after co-cultivation with AGS cells were counted (Fig. 2d). Here, we found that transfection of Wnt3 in AGS cells caused a substantial increase in HFE-145 cell number (Fig. 2d). Consistently, blockage of cytoneme formation in AGS cells and Wnt3 overexpressing AGS cells by IRSp53^4K^ co-transfection reduced HFE-145 cell number significantly. Next, we confirmed the observed alteration in cell numbers is based on proliferation by performing a BrdU incorporation assay to quantify actively proliferating cells with newly synthesized DNA (Fig.2e). First, we over-expressed Wnt3 in AGS cells and found a significant increase in the percentage of BrdU-positive cells (Fig. 2e, f). Consistently, we observed that blockage of cytoneme formation by over-expressing IRSp53^4K^ caused a reduction in proliferating cells. Finally, we showed that co-expression of Wnt3 together with IRSp53^4K^ led to a significant downregulation of actively proliferating cells. Our data provide the first evidence that Wnt3 localises to cytonemes and cytonemal-delivered Wnt3 can induce paracrine Wnt/β-catenin transduction cascade to promote cell proliferation in gastric cancer.

### Flot2 expression level correlates with cytonemal phenotypes in gastric cancer cells

To compare the molecular mechanism concerning the cytoneme-mediated Wnt3 dissemination in gastric epithelium cells and GC cells, we investigated the function of Flot2, a scaffolding protein, which enhances the generation of filopodia-like structures in various cancer cell lines (Neumann-Giesen *et al*., 2004) and is highly expressed in GC (Cao *et al*., 2013; Zhu *et al*., 2013). We found that AGS cells express 2.1-fold Flot2 mRNA and 2.7-fold higher levels of Flot2 protein than HFE-145 (Fig. 3a). Of all the GC cell lines, AGS cells express the highest Flot2 mRNA and protein levels (Suppl. Data Fig. 1c), which coincides with an increased number of filopodia (Fig. 1b). We next asked whether Flot2 is involved in the generation of filopodia in gastric epithelial cells as well as in gastric cancer cells. We found that Flot2 can induce a significant increase in the number and length of filopodia in HFE-145, similar to the ones observed in AGS cells (Fig. 3b-d). Expression of Flot2 in AGS does not further increase the average filopodia length due to promoting an increase in both longer and shorter filopodia, which averages a similar length to WT. Next, we blocked Flot2 function by using the dominant-negative Flot2 mutant ΔN-Flot2-GFP, which lacks its N-terminus and cannot localise to the membrane (Neumann-Giesen *et al*., 2004) and by an siRNA-mediated knock-down of Flot2. Consistently, transfection of ΔN-Flot2-GFP or Flot2 siRNA significantly decreased the filopodia length and number in both cell types (Fig. 3b-d). Control AGS cells as well as AGS and HFE-145 cells transfected with Flot2 display about 20% of filopodia with a length over 10µm (Fig. 3e). Concordantly, a reduction of Flot2 function led to an increase of short filopodia with a maximum length of 4µm. Additionally, cells with perturbed Flot2 function have more lamellipodia, suggesting a change in actin cytoskeletal dynamics (Supp. Fig. 2a). Thus, we conclude Flot2 enhances the formation and elongation of filopodia and thus might explain the differences of the cytonemal phenotype in the Flot2 low-expressing HFE-145 cells versus the GC cells displaying high levels of Flot2 expression.

**Figure 3.**
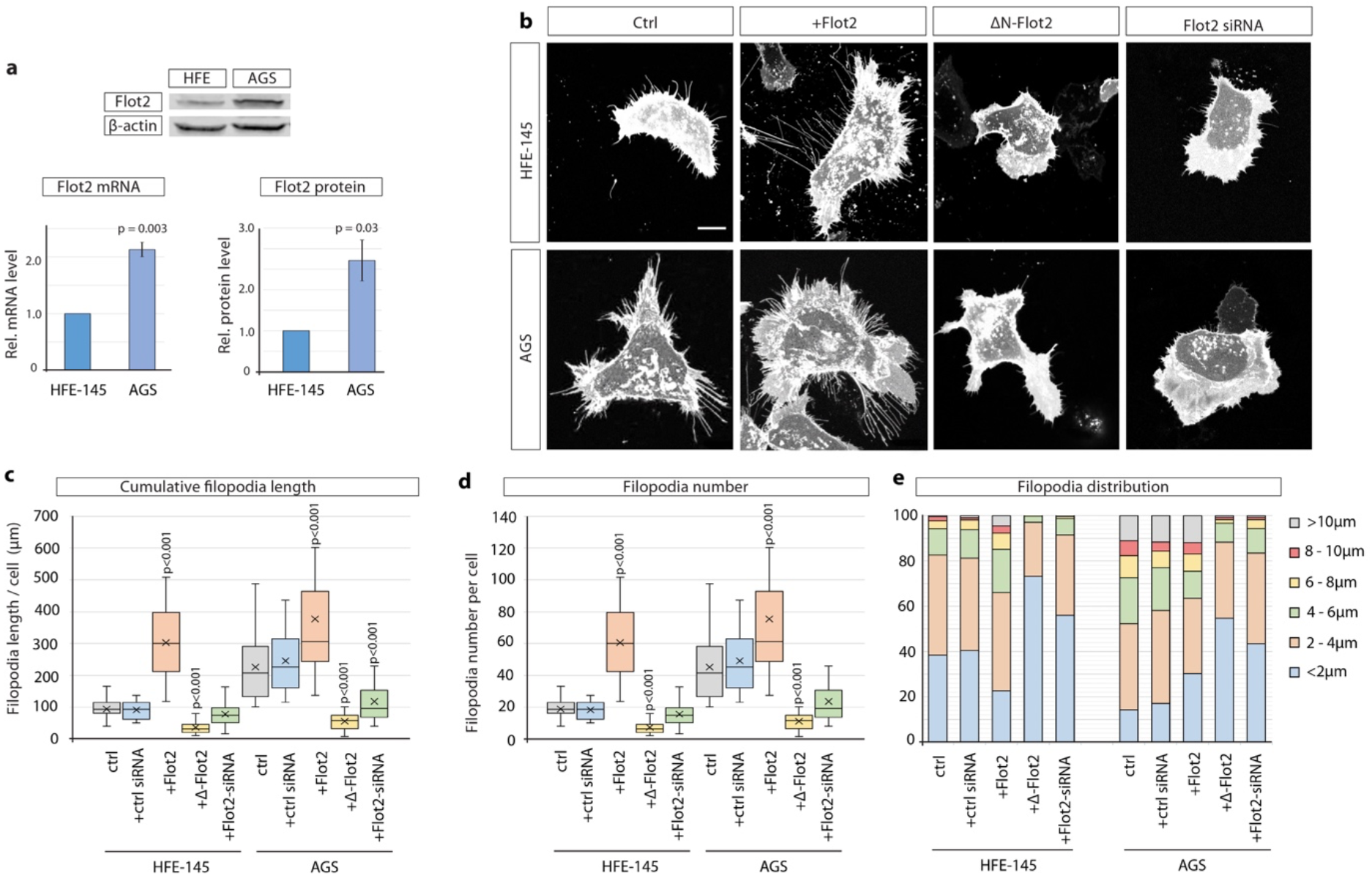
Flotillin-2 is over-expressed and promotes filopodia formation and elongation in gastric cancer cells. **a,** Flot2 protein levels in HFE-145 and AGS cells as quantified by Western Blot after normalising to beta-actin levels (n = 3) and by RT-qPCR after normalising to GAPDH as housekeeping gene (n = 4). Relative protein and mRNA levels are compared to HFE-145. Error bars represent SEM. Significance calculated by student’s t-test. **b,** Representative images of HFE and AGS cells expressing membrane-mCherry and indicated Flotillin-2 (Flot2) constructs or siRNA after 48hrs. Scale bars 10µm. **c-d,** Filopodia quantifications of HFE and AGS cells transfected with indicated Flot2 plasmids or siRNA. Significance calculated by Student’s t-test with Bonferroni correction for multiple comparisons. Average cumulative filopodia length (**c**), average filopodia number per cell (**d**). (n per condition (HFE) = 22, 19, 25, 23, 24). (n per condition (AGS) = 25, 21, 25, 25, 25; n = number of cells measured). **e,** Distribution of filopodia, categorized by length, as a percentage of total filopodia per HFE or AGS cell 48hrs post-transfection with indicated Flot2 plasmids or siRNA.

### Flot2 indicates Wnt3 cytonemes

Next, we investigated if Flot2 specifically regulates Wnt3 cytonemes, rather than just filopodia. Therefore, we investigated whether Flot2 and Wnt3 are co-localising on cytonemes. In AGS cells, we found that Flot2 displays punctate staining at the plasma membrane with noticeable staining on filopodia (Fig. 4a). Next, we investigated whether these filopodia contain both, Wnt3 and Flot2. After double IHC analysis, we found that Flot2 and Wnt3 are co-localising on Wnt3 cytonemes (Fig. 4b). Similar to endogenous Flot2, Flot2-GFP localised along the length and to tips of filopodia (Fig. 4c). Intriguingly, we detected that Flot2-GFP co-localises with Wnt3-mCherry on cytonemes (Fig. 4d), similar to the endogenous protein. We further observed that Flot2-GFP and Wnt3-mCherry co-localise strongly at the cytoneme contact sites (Fig. 4d). From this data, we hypothesised that Flot2 is a marker of Wnt3 cytonemes in GC cells.

**Figure 4.**
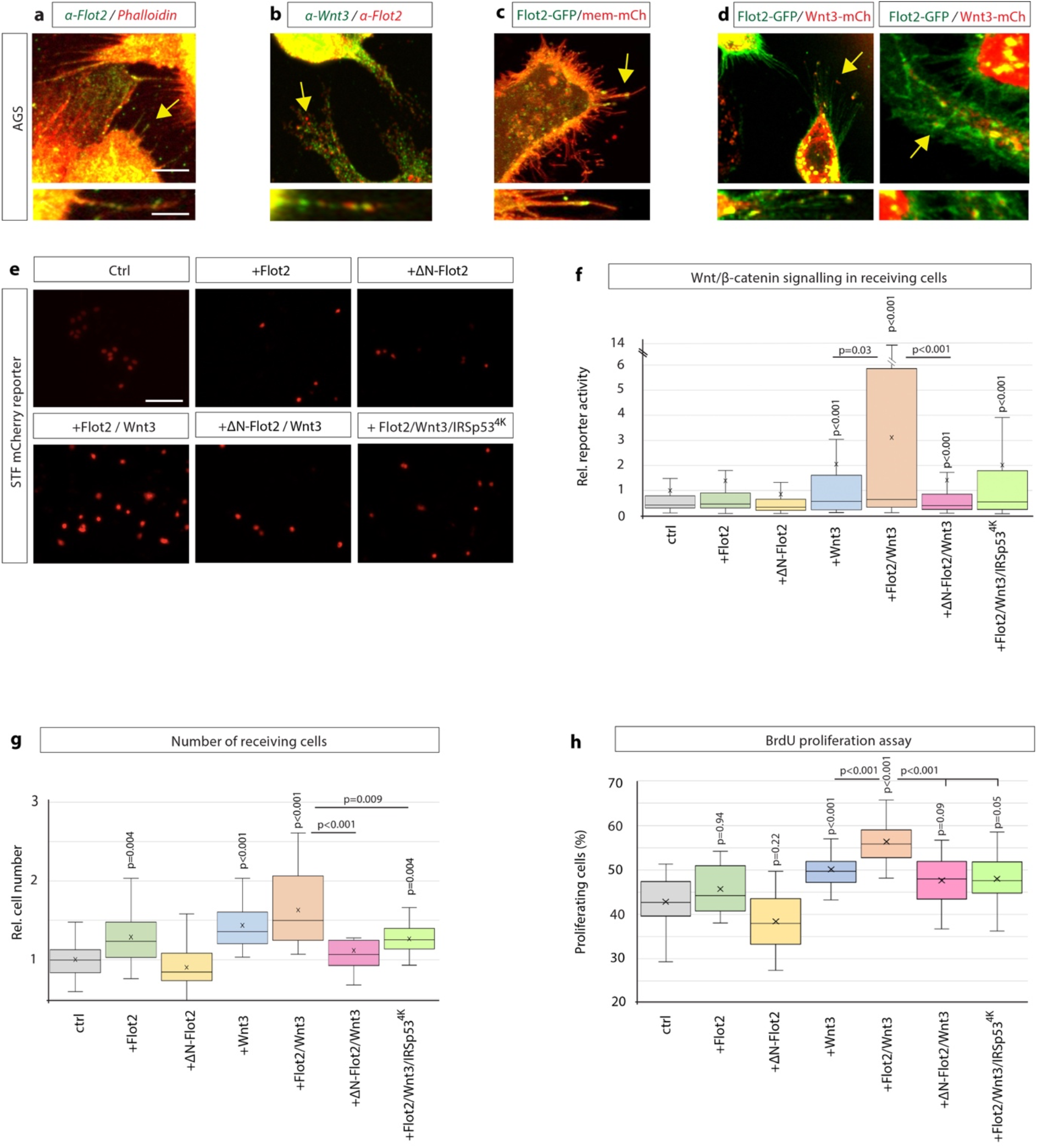
Flotillin-2 marks Wnt3 cytonemes and regulates paracrine Wnt/β-catenin signalling and proliferation. **a,** IHC analysis showing endogenous localisation of Flot2 (green) in AGS cells. TRITC phalloidin was used to visualise actin. Arrows indicate localisation of Flot2 to filopodia. Scale bars 5µm. High-magnification images indicate an example of a Flot2-bearing cytonemes. Scale bars 2.5µm. **b**, IHC analysis show that Flot2 co-localises with Wnt3 on cytonemes. **c,** Confocal images showing the sub-cellular localisation of Flot2-GFP in AGS cells. Arrows indicate localisation of Flot2-GFP on cytonemes. **d,** Confocal images highlighting co-localisation of Flot2-GFP and Wnt3-mCh on cytonemes in AGS cells (arrows). Flot2-GFP and Wnt3-mCherry also cluster and co-localising at a cytoneme contact point (arrow). **e,** Representative images of STF reporter fluorescence for indicated conditions. Scale bar 100uM. **f,** Relative quantification of STF mCherry reporter fluorescence in HFE cells co-cultivated with AGS cells expressing indicated constructs. Quantifications relative to AGS control. (n per condition = 322, 443, 403, 258, 336, 306, 297; n = number of nuclei measured). **g,** Relative number of HFE cells per image after co-cultivation with AGS cells expressing indicated constructs. Significance calculated by student’s t-test with Bonferroni correction for multiple comparisons. (n per condition = 28, 27, 26, 27, 22, 24, 15; n = number of images). **h,** Quantification of BrdU-stained cells as a percentage of the population, after co-cultivation of AGS and HFE cells, as described in Fig. 2a. Significance calculated by student’s t-test with Bonferroni correction for multiple comparisons. (n per condition = 20; n = number of images).

### Paracrine Wnt/β-catenin signal activation and proliferation are enhanced by Flot2

To investigate a functional relationship between Wnt3 and Flot2, we assessed whether Flot2 function has an impact on paracrine Wnt signalling via cytonemes. First, we performed a co-cultivation assay of AGS cells and HFE-145 cells as done previously (Fig. 2a). We observed that expression of Flot2 in the AGS cells slightly increased the expression of the STF mCherry reporter in the neighbouring HFE-145 cells (Fig. 4e). Consistently, ΔN-Flot2-GFP reduced paracrine Wnt signal activation in HFE-145. Strikingly, co-expression of Flot2 and Wnt3 significantly enhanced the STF reporter expression and thus, paracrine Wnt signal activation. This increase is reduced upon co-expression of Wnt3 with ΔN-Flot2-GFP. Even more striking, the expression of the cytoneme antagonist IRSp53^4K^ led to a significant reduction of paracrine Wnt signalling activation in Wnt3 and Flot2 expressing cells. Our results suggest that Flot2 significantly increases the number of Wnt3 cytonemes (Fig. 3, 4) and this increase led to a massive upregulation of paracrine Wnt signal activation (Fig. 4e).

Next, we determined the impact of these changes in Wnt signal activation on cell number in Wnt-receiving HFE-145 cells (Fig. 4g). We found that Flot2 expression caused a significant increase in HFE-145 cell number, whilst ΔN-Flot2 caused a decrease. Following a similar trend to the STF reporter, co-expression of Flot2 with Wnt3 resulted in the greatest increase, whereas the Wnt3-induced signalling was attenuated by co-expression with ΔN-Flot2-GFP. Similarly, IRSp53^4K^ attenuated increases seen in the presence of Wnt3/Flot2, suggesting Flot2 has a significant impact on proliferation in a cytoneme-dependent manner. The effects of Flot2-Wnt3 signalling on proliferation were confirmed by a BrdU based cell proliferation assay. We found that co-expression of Flot2 and Wnt3 produced the greatest increase in BrdU incorporation into newly synthesized DNA of actively proliferating cells. (Fig 4h). Consistently, IRSp53^4K^ reduced BrdU labelling and thus proliferation. Our data further suggest Flot2-dependent cytonemes are a key transport mechanism for Wnt3 in gastric cancer, regulating paracrine Wnt signalling and GC cell proliferation.

### Intracellular transport and membrane localisation of Ror2 is influenced by Flot2

Next, we addressed the underlying mechanism for Flot2-mediated cytoneme formation. Previously, the Wnt co-receptor Ror2 has been shown to be critical for Wnt-mediated cytoneme formation and signalling (Mattes *et al*., 2018; Brunt *et al*., 2021). Since Ror2 has been shown to localise to lipid rafts (Sammar *et al.,* 2009), we hypothesised that Flot2 could be interacting with Ror2 in promoting cytoneme formation. To test this hypothesis, we analysed first the intracellular distribution and found striking co-localisation of Ror2 and Flot2, particularly at clusters at the plasma membrane (Fig. 5a), most likely Flot2 positive microdomains. Overexpression of tagged version of Flot2 and Ror2 led to a similar co-localisation at the membrane, and, consequently, the formation of long cytonemes (Fig. 5b). Consistently, expression of a dominant negative form of Flot2 resulted in a strong reduction of cytonemes (Fig. 5c). In parallel, we found that reduced Flot2 function led to a loss of Ror2 membrane localisation and, subsequently, Ror2 accumulated in the perinuclear region (Fig. 5c). We observed a similar phenotype after knock-down of Flot2 - a reduction of cytonemes and a removal of Ror2 from the membrane (Fig. 5d). These experiments suggest that Ror2 requires Flot2 for proper membrane localisation. To further assess the change in subcellular localisation, we mapped the localisation of Ror2 with regard to a set of organelle markers (Fig. 5e) and found that upon siRNA-mediated Flot2 knockdown, Ror2-mCherry localisation significantly increases in the Golgi apparatus without a noticeable change in the other organelles investigated (Fig. 5f). This data set suggests Flot2 is required for Ror2 trafficking between the Golgi and the membrane.

**Figure 5.**
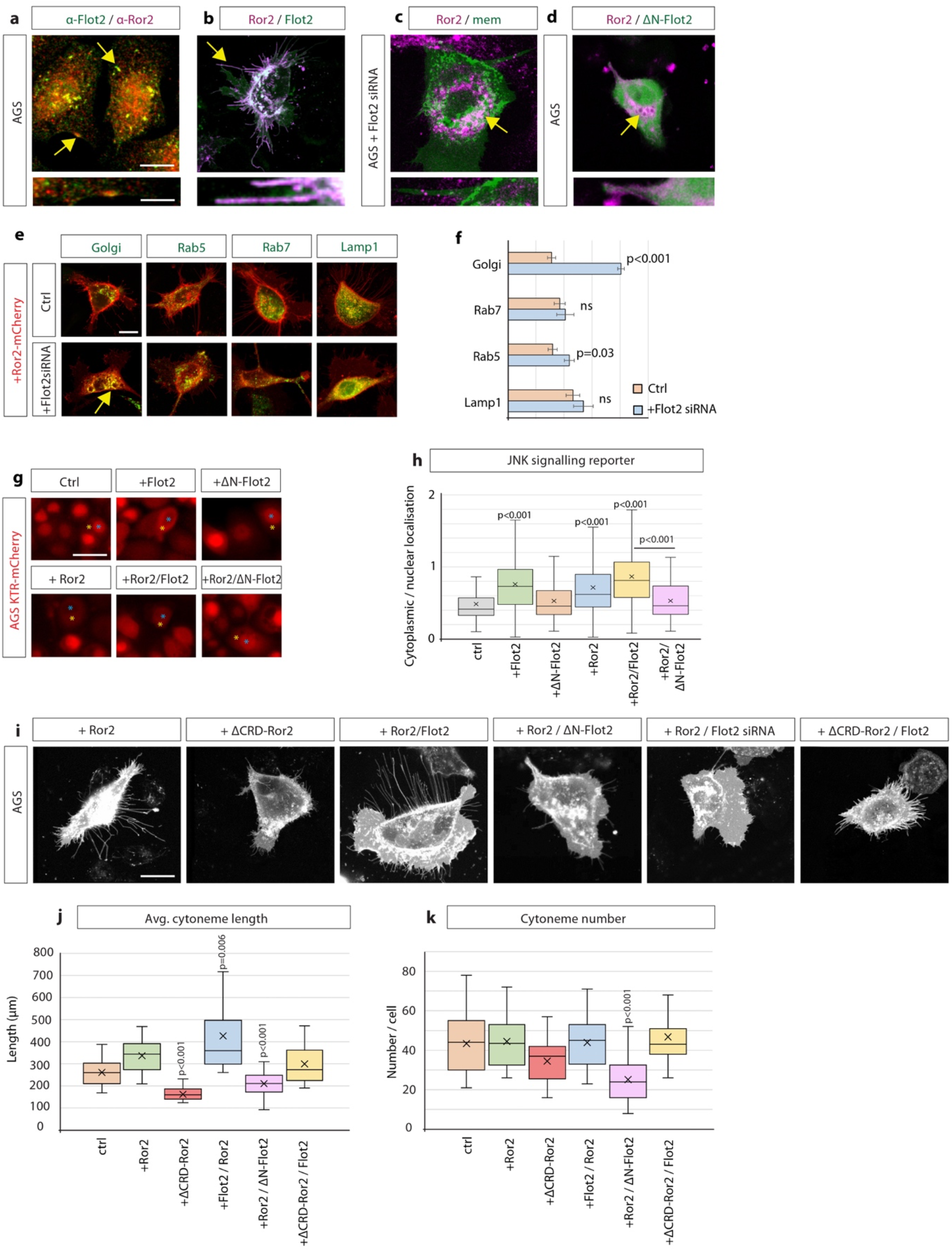
Flotillin-2 is required for Ror2 membrane localisation, Ror2/PCP signalling and Ror2-mediated cytoneme formation. **a,** IHC analysis of AGS cells stained for Ror2 (red) and Flot2 (green). Co-localisation of Flot2 and Ror2 at the membrane highlighted by arrows. Scale bars represent 10µm and in high-magnification images 2.5µm (right). **b-d**, Confocal live cell imaging of AGS cells expressing Ror2-BFP with Flot2-GFP (**b**), Flot2 siRNA (**c**) or ΔN-Flot2-GFP (**d**) and memCherry. **e,** Live confocal images of AGS cells expressing Ror2-mCherry and indicated organelle markers +/-Flot2 siRNA. Arrows highlight co-localisation of Ror2-mCherry and mTurq2-Golgi. Scale bar 10µm **f,** Quantification of co-localisation of Ror2-mCherry with indicated markers, assessed by Pearson’s Correlation Co-efficient (PCC). Significance calculated by Student’s t-test. (n per condition (WT) = 7, 10, 8, 10) (n per condition (Flot2 siRNA) = 7, 6, 7, 8). **g,** Representative images of AGS cells stably expressing the JNK-KTR-mCherry reporter and indicated constructs after 48 hrs. Yellow asterisks marks cytoplasmic localisation, blue asterisks mark a nuclear localisation. Scale bar 20µm. **h,** Quantification of the JNK-KTR-mCherry reporter. Nuclear and cytoplasmic fluorescence of cells were measured and the cytoplasmic : nuclear ratio calculated. Significance calculated by one-way ANOVA with Bonferroni correction for multiple comparisons. (n per condition = 136, 109, 74, 109, 82, 79). **i,** Representative confocal images of AGS cells expressing memCherry and indicated constructs for 48 hrs. Scale bars 10µm. **j, k,** Quantification of cytoneme length and number of AGS cells transfected with constructs indicated in (**i**). Significance calculated by Student’s t-test with Bonferroni correction for multiple comparisons. (n per condition = 25, 22, 21, 23, 25, 21; n = number of cells).

### Ror2/PCP signalling is regulated by Flot2

Based on the observation that Flot2 is required for proper membrane localisation of Ror2, we hypothesized that Ror2 mislocalisation would also impact on Ror2-mediated cytoneme formation (Mattes *et al.,* 2018). During cytoneme outgrowth, Ror2 needs to activate the Wnt/PCP pathway, including downstream activation of JNK signalling, to induce polymerisation of the cytonemal actin cytoskeleton (Martinez *et al*., 2015, Brunt *et al.,* 2021). Since Flot2 is required for Ror2 membrane localisation, we investigated if Flot2 function can be linked to Ror2/PCP signalling. Therefore, we generated an AGS cell line stably expressing a reporter of JNK signalling, the JNK kinase translocation reporter (JNK KTR-mCherry) (Regot *et al*., 2014; Miura *et al*., 2018; Brunt *et al*., 2021). JNK-KTR-mCherry localises to the nucleus (N) in its dephosphorylated state (low JNK activity). Upon activation by phosphorylation, it shuttles to the cytoplasm (C, high JNK activity). The C:N ratio can then be calculated as an indicator of JNK signalling. AGS cells display a low basal levels of JNK activity (Fig. 5g, h). Expression of either Flot2 or Ror2 significantly increases JNK signalling, and co-expression of Flot2 / Ror2 further enhances JNK signalling activation, suggesting a synergistic effect. In accordance with a synergistic function, we demonstrated that blockage of Flot2 function in Ror2 positive AGS cells by co-expression of a ΔN-Flot2, can reduced PCP/JNK signalling significantly. This data set suggests that Flot2 is required for Ror2-induced JNK signalling. In parallel, we observed that Flot2 represses autocrine β-catenin signalling pathway, which is consistent with previous observations suggesting a mutually repressive nature of the Wnt/PCP and Wnt/β-catenin pathways (Supp. Fig. 3a, b) (van Amerongen and Nusse, 2009; Niehrs, 2012). Together, these data imply that Flot2 is required for Ror2-induced JNK signalling but inhibits canonical signalling in the Wnt-producing cell.

### Ror2 and Flot2 are required for cytoneme formation

Finally, we wanted to decipher the combined role of Flot2 together with Ror2 in cytoneme induction. For this reason, AGS cells were transfected with WT or mutant Flot2 and Ror2 constructs. membrane-mCherry expression was used to visualise and quantify cytonemes (Fig. 5i-k). Expression of Ror2 alone increased the average cytoneme length whilst having no significant effect on cytoneme number (Fig. 5j,k). However, co-expression of Flot2 with Ror2 enhanced this phenotype; increasing cytoneme length. Consistently, blockage of Flot2 function upon expression of the dominant-negative ΔN-Flot2-GFP decreased the number and length of cytonemes in Ror2 expressing AGS cells (Fig. 5j,k). To test whether Flot2 requires Ror2 for cytoneme formation, a mutant construct of Ror2 missing the Wnt-interacting CRD domain (ΔCRD-Ror2) was used, which cannot bind to Wnt ligands and thus transduce a Wnt signal. When expressed in AGS cells, ΔCRD-Ror2 significantly reduced the average length and number of cytonemes. We found that this phenotype can be partially rescued by the expression of Flot2, which moderately restores the average cytoneme length and number, suggesting that Flot2 acts downstream of Ror2-inducing cytonemes. Taken together, these results show that Flot2 is required for the membrane localisation of Ror2. Furthermore, Flot2 facilitates Ror2-mediated Wnt/PCP signalling and, consequently, the promotion of Wnt cytonemes.

### Wnt8a cytonemes in zebrafish development require Flot2 function

To evaluate whether the role of Flot2 in formation of Wnt cytonemes is conserved, we addressed Flot2 function during zebrafish development. Previous work from our lab has shown that cytonemes are essential for the dissemination of Wnt8a in zebrafish embryogenesis (Stanganello *et al*., 2015) and that these cytonemes can be regulated by the Ror2/PCP pathway (Mattes *et al*., 2018; Brunt *et al*., 2021). First, we mapped the subcellular localisation of Flot2 in zebrafish epiblast cells and found that Flot2-GFP displayed strong membrane localisation and was localised to filopodia, including their tips (Fig. 6a). Flot2-GFP could also be seen clustering together with Wnt8a-mCherry on cytonemes (Fig. 6b), analogous to the co-localisation seen with Wnt3 in GC cells. Next, we altered Flot2 function during zebrafish embryogenesis and observed that the expression of Flot2 significantly increased the average cytoneme length and cumulative length, whilst having no significant effect on cytoneme number (Fig. 6c, d; Supp. Fig. 3c). Consistently, the dominant-negative ΔN-Flot2-GFP attenuated the cytoneme number and cumulative length. This data confirms that Flot2 regulates cytoneme in the zebrafish embryo, concurrent with our findings in GC cells.

**Figure 6.**
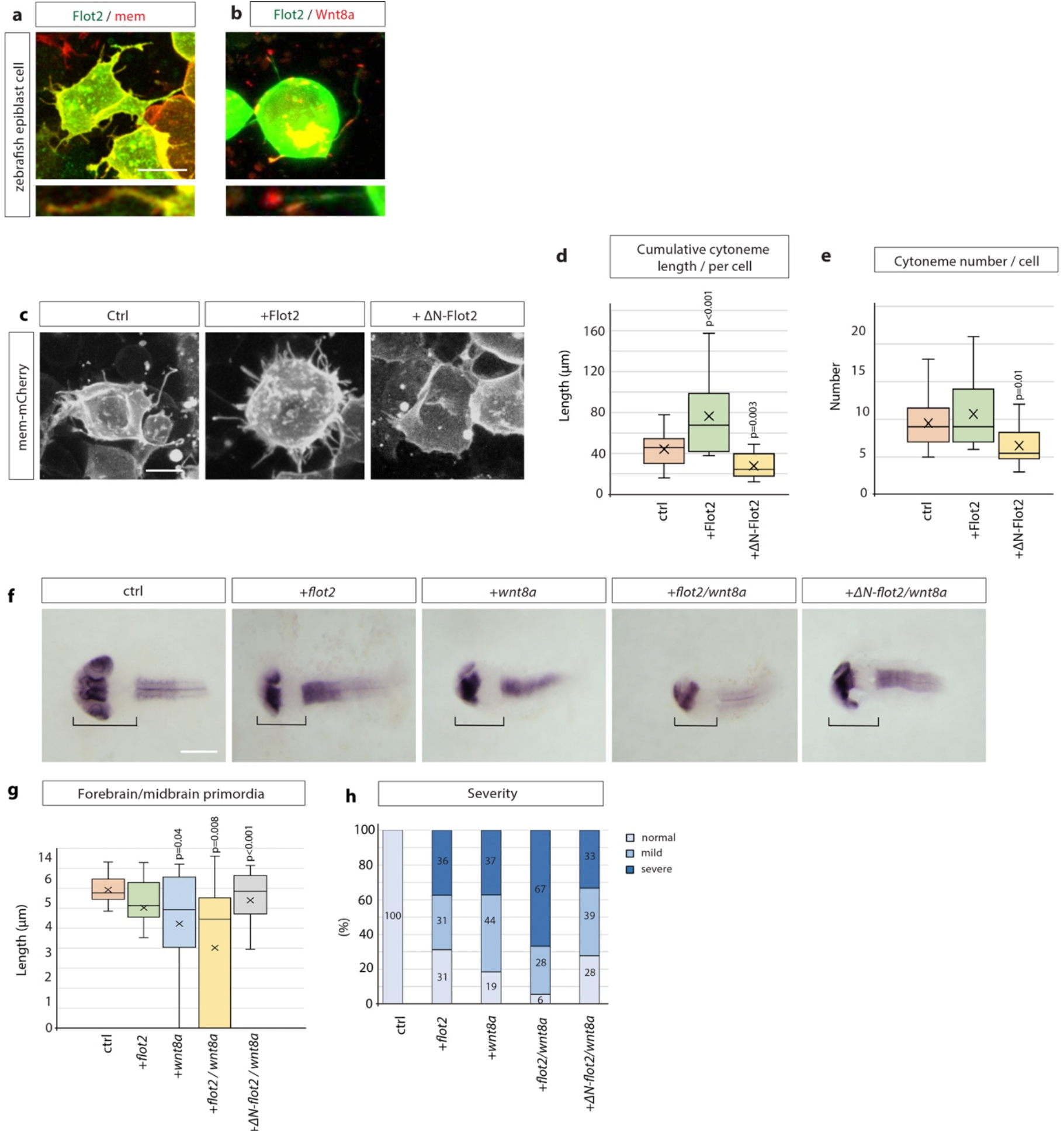
Flotillin-2 promotes cytoneme formation and Wnt8a signalling in zebrafish development. **a, b,** Confocal images of zebrafish epiblast cells injected with Flot2-GFP and memCherry DNA (100ng/µl) (**a**) and Flot2-GFP and Wnt8a-mCherry (100ng/µl) and imaged at 8 hpf. Scale bars represent 10µm **c,** Representative images of zebrafish epiblast cells injected with memCherry indicated constructs. Scale bar 10µm. **d, e**, Quantification of filopodia from epiblast cells injected in **c**. Significance calculated by Student’s t-test. (n per condition = 17, 20, 14). **f,** *In situ* hybridisation against *pax6a* in zebrafish embryos at 30hpf after microinjection of 100ng/µl of indicated DNA constructs and imaged. Scale bar represents 100µm. **g,** Quantification of forebrain and midbrain primordia length in zebrafish embryos injected as in (**f**). Significance calculated by Student’s t-test. (n per condition = 23, 16, 27, 18, 18; n = number of embryos). **h,** Qualitative analysis of phenotype severity in zebrafish embryos injected as indicated in (**f**). Phenotypes classified into the categories normal, mild and severe. Numbers in bars represent percentages of total embryos.

Next, we addressed the consequences of Flot2 function on the formation of the Wnt/β-catenin signalling gradient in zebrafish embryogenesis. Wnt8a is a key Wnt morphogen in determining the positions of the boundaries of the brain anlage, and alteration of Wnt8a cytonemes perturbs brain anlage patterning (Mattes *et al*., 2018; Brunt *et al*., 2021). Therefore, we altered Wnt expression levels together with Flot2 levels during neural plate patterning and found that the expression of low levels of *wnt8a* mRNA results into an anterior shift of the boundaries of the brain anlage and reduces the combined forebrain (FB) and midbrain (MB) length (Fig. 6f, g). Such a posteriorisation was also observed upon injection of *flot2* mRNA. An even more pronounced shift of the position of brain anlage boundaries occurred upon co-injection of *flot2* and *wnt8a* mRNA, suggesting a synergistic effect of Flot2 and Wnt8a during neural plate patterning. Consistently, blockage of Flot2 function by expression of the dominant-negative ΔN-Flot2-GFP attenuated the alteration in patterning. As well as perturbing the length of the primordia of the FB/MB, alteration of Wnt8a signalling can cause defects in the early zebrafish embryo, which can include complete loss of the FB primordium and under-developed or missing eyes. We categorised the developmental defects and found that embryos co-expressing Flot2 and Wnt8a significantly increased the number of embryos with severe defects (Fig. 6h).

Together, these data suggest Flot2 can enhance the length and number of Wnt8a cytonemes, and thus alter the Wnt8a signalling gradient, leading to a posteriorisation of the zebrafish brain anlage. These findings are concurrent with our results in GC cells showing Flot2 enhances paracrine Wnt signal activation. In both systems, Flot2 also co-localises with the Wnt ligands, Wnt3 and Wnt8a, and promotes cytoneme formation, which suggests a conserved role for Flot2 in promoting cytoneme-mediated Wnt dissemination in vertebrate tissue.

## Discussion

Wnt/β-catenin pathway activity, in addition to other stemness signals, is required to maintain the gastrointestinal crypt microenvironment (Sato *et al*., 2011). Wnt3 is an essential short-range signal at the crypt base (Farin, Van Es and Clevers, 2012), which does not freely diffuse (Farin *et al*., 2016). Similarly, our study of gastric epithelial cells shows that Wnt3 remains membrane-tethered during transport and signalling (Fig. 1). In support of these observations, experiments in *Drosophila* suggest that Wg is similarly not freely diffusible as flies with a membrane tethered form of Wg develop largely normal (Alexandre, Baena-Lopez and Vincent, 2014). Here, we provide evidence that gastric epithelial cells can load Wnt3 on cytonemes for intercellular distribution (Fig. 1). In gastrointestinal adenomas and differentiated carcinomas, Wnt3 and Wnt cargo receptor Evi/Wntless are significantly increased and crucial for cancer cell proliferation and colony formation (Voloshanenko *et al*., 2013). Furthermore, the Wnt receptor Fzd7, which can bind Wnt3, transmits oncogenic Wnt signalling to drive the proliferation of gastric tumours *in vivo* (Flanagan *et al*., 2019). In conclusion, elevated levels of Wnt3 are essential for quickly proliferating gastrointestinal cancers, however, it is unclear how this hydrophobic ligand is disseminated within a tumour’s context. In many cancer cells, the number of filopodia is increased (Machesky and Lia, 2010) and LGR5^+^ intestinal stem cells form long cytonemes (Snyder *et al*., 2015). Our data suggest that the cancer stem cell like AGS cells form a similarly high number of long filopodia and we assign a fresh function to these filopodia (Fig. 1). We propose that enhanced Wnt3 expression and an increase of cytonemes lead to augmented Wnt3 dissemination and signalling within GC cells, a prerequisite for adenoma formation. In support of our hypothesis, in *Drosophila,* cytoneme-mediated FGFR signalling is similarly required for tumour growth and malignancy (Fereres *et al*., 2019). However, the underlying molecular mechanism increasing cytoneme emergence in cancers is unclear.

Flotillins are upregulated in many tumours (Gauthier-Rouvière *et al*., 2020 Zhu *et al*., 2013). Our data suggest that Flot2 expression correlates with enhanced cytoneme formation in gastric cancer. These results concur with previous studies demonstrating the ability of Flot2 to enhance filopodial phenotypes (Hazarika *et al*., 1999; Neumann-Giesen *et al*., 2004). We further show that increased Flot2-function leads to a further Wnt ligand dissemination in gastric cancer cells as well as in zebrafish embryogenesis. These observations are in concert with data in *Drosophila* showing that Flot2 promotes long-range Wg signalling in the wing imaginal disc (Katanaev *et al*., 2008). Consequently, blockage of Flot2 function reduces paracrine Wnt signalling and cell proliferation significantly – despite the high levels of Wnt3 expressed in GC cells (Fig. 4). We, therefore, speculate that GC cells require not only increased Wnt3 expression levels for tumour maintenance but that increased Flot2 expression allows more effective ligand dissemination in the tumour tissue.

Previously, Wnt/PCP signalling has been recognised as a key regulator of Wnt cytoneme emergence in vertebrates (Mattes *et al*., 2018; Brunt *et al*., 2021), where the receptor tyrosine kinase-like orphan receptor Ror2 has been implicated in activating this signalling pathway (Ho *et al*., 2012). Furthermore, autocrine Ror2/PCP signalling leads to the induction of Wnt cytonemes in zebrafish gastrula and mouse myofibroblasts (Mattes *et al*., 2018). Here, we provide evidence that Ror2 is similarly required for Wnt cytoneme formation in GC via the activation of Wnt/PCP/JNK signalling. However, we found that GC cells display more and longer filopodia despite sequencing data indicating that Ror2 expression is reduced in GC tissue (Li *et al.,* 2014). We, therefore, suggest that the low levels of Ror2 seen in GC are functionally compensated by high levels of Flot2, which works synergistically to induce long cytonemes.

Surprisingly, we also found that Flot2 function is a prerequisite for Ror2-mediated Wnt/PCP signalling, as blocking Flot2 function reduces Ror2 membrane localisation, signalling and cytoneme induction. In accordance with our findings, it has been suggested that Flot2 can influence receptor tyrosine kinase (RTK) signalling by providing a signalling microdomain at the plasma membrane (Banning *et al*., 2014). Furthermore, there is good evidence that Flotillins influence EGF, FGF, and MAPK signalling, however, Flotillins have contradictory effects depending on the experimental setting and the tissue context and a unifying explanation is missing to date (Amaddii *et al*., 2012; Banning *et al*., 2014). We hypothesise that Flot2 - in addition to its function in organizing microdomains at the membrane - is a key regulator of the intracellular protein transport. Thus, we suggest that Flot2 function is a prerequisite for activating the Wnt/PCP dependent cytonemes; however, further functional roles for Flot2 might exist and might also be modulated in a cell-type-specific manner.

As a mechanistic explanation, we suggest that Flot2 regulates intracellular trafficking of the Wnt/PCP co-receptor and RTK Ror2 from the Golgi apparatus to the plasma membrane for cytoneme induction. On one hand, this is consistent with reports highlighting Flot2 as a regulator of membrane invagination and trafficking between endocytic compartments (Frick *et al*., 2007). In particular, it has been observed that cargo, including Flot2 itself, accumulate in the Golgi complex of HeLa, Jurkat and PC12 cells when Flot2 function is obstructed (Langhorst *et al*., 2008). Consistently, we found that ectopic Flot2 expression leads to an effective transport from Ror2 to the plasma membrane and, thus, to enhanced Wnt3 cytoneme formation in the source cells (Fig. 5). Consequently, paracrine Wnt/β-catenin signalling is facilitated in neighbouring receiving cells. In accordance, we found that blockage of Flot2-induced cytonemes - without interfering with Wnt3 expression and secretion - can significantly reduce paracrine Wnt activation (Fig. 4). This suggests that Wnt3 uses mainly Flot2/Ror2 dependent cytonemes for intercellular transport. Whether the canonical Wnt3 is a transient passenger on these cytonemes, directly binding to Ror2 or binding to Evi/Wntless, remains to be answered. However, there is indeed some evidence that Wnt3 can bind to Ror2 (Billiard *et al*., 2005). Furthermore, Wnt8a is a further example of a Wnt/β-catenin ligand interacting with not only multiple Frizzled receptors but also both Ror2 and Lrp6 co-receptors (Mattes *et al*., 2018). Clarifying whether Wnt3 displays similar capabilities will aid our molecular understanding of its cytoneme localisation.

It has been suggested that elevated Wnt/β-catenin signalling activity correlates with stem cell signatures in cancer (Reya and Clevers, 2005). Therefore, it is tempting to speculate that gastric cancer cells generate a ‘niche’-like environment that allows them to maintain stemness. A crucial factor for maintaining the gastric cancer stem cell niche is elevated Wnt3 signalling. We show that gastric cancer stem cells can form a cytonemal network allowing a fast and precise exchange of Wnt3. As an underlying mechanism, we, therefore, suggest that Flot2 is crucial in regulating cytoneme emergence in the niche to facilitate long-range Wnt3 transport. Indeed, the expression levels of both Wnt3 and Flot2 correlate with the degree of intestinal tumour progression and with poor prognosis (Merlos-Suárez *et al*., 2011; Liu *et al*., 2018). We propose that despite the presence of pathway-activating mutations in APC or β-catenin, the survival of intestinal cancer cells remains dependent on Wnt ligand distribution on Flot2-controlled cytonemes. Thus, besides interfering with the Wnt signalling pathway directly, regulating Flot2-mediated Wnt cytonemes in the cancer stem cell niche could provide a novel strategy for combatting Wnt-related cancers in the future.

## Acknowledgements

The authors are funded by the Medical Research Council (MRC)/UKRI through a studentship for D.R. (MR/N0137941/1 for the GW4 BIOMED DTP) awarded to the Universities of Bath, Bristol, Cardiff and Exeter, and a research grant (MR/S007970/1) awarded to S.S. We would like to thank Ritva Tikkanen for generously providing the Flotillin-2 constructs used in this study. We would like to thank Sahin Deniz (Istanbul) and Duane T. Smoot (Nashville) for generating the HFE-145 gastric epithelial cell line. Finally, we would also like to thank Chengting Zhang and Lucy Brunt for technical help, and the entire Scholpp lab for their comments on the manuscript.

## Author contributions

D.R., T.J.P., and S.S. designed experiments and wrote the manuscript. D.R. performed all experiments and data analysis. S.R. generated some of the DNA plasmids used and the stable JNK8-AGS cell line. H.A. provided the HFE cell line used in this study.

## Methods and Materials

### Plasmids and Antibodies

The following plasmids were used in transfections and/or microinjections: pCS2+ membrane-mCherry (Scholpp et al., 2009), pCAG-mGFP membrane-bound GFP (Addgene 14757), pEGFP-N1 Flot2-GFP (Neumann-Giesen et al., 2004), pEGFP-N1 ΔN-Flot2-GFP (Neumann-Giesen et al., 2004), 7xTRE Super TOPFlash-NLS-mCherry (Moro et al., 2012), JNK KTR-mCherry (Regot et al., 2014), pCS2+ LifeAct-GFP, pCS2+ Rab5-GFP (Jim Smith Group), Rab7-eGFP (from Rüdiger Rudolf), pEGFP-N1 LAMP1-mTurq2 (Addgene #98828), IRSp534K-mCherry/GFP (Stanganello et al., 2015), pCS2+ Wnt8a-mCherry (Stanganello et al., 2015), pCS2+ Ror2-mCherry (cloned in with ClaI and XbaI), pEGFP-N3 mRor2-ΔCRD-GFP (Minami et al., 2010), pCS2+ Ror2-eBFP2 (cloned by inserting eBFP2 into pCS2+ Ror2 plasmid using XbaI and SnaBI), pCS2+ Wnt3-mCherry (cloned from Addgene plasmid pcDNA3.2-Wnt3 (#35909) into pCS2+-mCherry vector using ClaI and XbaI).

The following primary antibodies were used for immunofluorescence and/or Western blots: anti-Wnt3 (Abcam 116222), anti-Flotillin-2 (Abcam 96507), anti-Flotillin-2 (Abcam ab113661), anti-Ror2 (Santa Cruz H-1), anti-beta-actin (Proteintech® 60008-1-Ig), anti-Myosin-X (Santa Cruz C-1), anti-Evi (EMD Milipore YJ5). The following AlexaFluor® (Thermofisher) secondary antibodies were used for immunofluorescence: Goat anti-rabbit 488 (Abcam 150077), Goat anti-rabbit 568 (Abcam 175471), Donkey anti-goat 647 (Abcam ab150135). For Western blotting, goat anti-mouse IRDye®800CW (Abcam 216772) and donkey anti-rabbit AlexaFluor®680 (Abcam 175772) were used.

### Cell Culture

The following cell lines, kindly authenticated and gifted from Dr. Toby Phesse (Cardiff University, UK), were used: gastric tubular adenocarcinoma cell line MKN7, derived from metastatic site (lymph nodes) and the gastric tubular adenocarcinoma cell line MKN28, derived from metastatic site (liver) and the primary gastric adenocarcinoma cell line AGS (Flanagan *et al*., 2019). As a control, the non-neoplastic epithelial cell line HFE-145, gifted from Dr. Hassan Ashktorab (Howard University, Washington, USA) was used. All cell lines were tested regularly for mycoplasma by endpoint PCR testing every 3 months and broth tests every 12 months.

AGS, MKN7 and MKN28 cells were maintained in RPMI-1640 (Sigma-Aldrich), and HFE-145 in DMEM (Thermofisher) media, all supplemented with 10% FBS and 1% Pen-Strep (Gibco). Cells were routinely passaged (with 0.05% EDTA-free trypsin (Thermofisher)) at ∼80% confluency.

Transient transfections of cells, either reverse or forward, were performed using FuGeneHD (Promega) according to manufacturer’s protocol (using 3:1 FuGene:DNA ratio). siRNA transfections were performed using Lipofectamine RNAiMAX (Thermofisher) according to manufacturer’s protocol, with a final siRNA concentration of 30 pmol. Control siRNA (MISSION® siRNA Universal Negative Control #1) and Flotillin-2 siRNA (Thermo Fisher 122408) were used.

### Zebrafish Maintenance and Husbandry

WIK wild-type zebrafish (Danio rerio) were maintained at 28°C and on a 14hr light/10hr dark cycle (Brand *et al.,* 2002). Zebrafish care and all experimental procedures were carried out in accordance with the European Communities Council Directive (2010/63/EU) and Animals Scientific Procedures Act (ASPA) 1986. In detail, adult zebrafish for breeding were kept and handled according to the ASPA animal care regulations and all embryo experiments were performed before 120 hrs post fertilization. Zebrafish experimental procedures were carried out under personal and project licenses granted by the UK Home Office under ASPA, and ethically approved by the Animal Welfare and Ethical Review Body at the University of Exeter.

### 7xTCF-NLS-mCherry TOPFlash Assays

For autocrine signalling (single cell population), 1×10^6^ HFE or AGS cells were reverse transfected with 7xTRE Super TOPFlash reporter plasmid along with indicated plasmids. Cells were incubated for 48 hrs before image acquisition on a Leica DMI6000 SD with a 20x objective.

For co-cultivation assays, 2×10^6^ HFE-145 cells were reverse transfected with 7xTRE Super TOPFlash plasmid (7xTCF-NLS-mCherry) and 2×10^6^ AGS cells with indicated plasmids in 6 well plates. After 24 hrs incubation, both cell types were trypsinised, counted and 1×10^6^ of each were co-cultivated in 6 well plates for a further 48 hrs before image acquisition.

For all assays, at least 5 images were taken in random locations for each biological repeat on a 20x objective. Fluorescence intensity of nuclei were measured on Fiji software. The number of fluorescent nuclei were also counted as a measure of proliferation.

### JNK KTR-mCherry Assay

AGS cells stably expressing JNK-KTR-mCherry (AGS-JNK8) were reverse transfected with indicated plasmids and incubated for 48 hrs. Cells were imaged on a Leica DMI6000 SD with a 20x objective. At least 5 images were taken in random locations for each biological repeat. Fluorescence intensity of each cell’s cytoplasm and nucleus was measured using Fiji software. After background subtraction, cytoplasmic:nuclear ratio (C:N) was calculated.

### Western Blotting

Cell lysates were collected from cell pellets by resuspending in 100uL of ice cold TNT lysis buffer (20mM Tris pH8.0, 200mM NaCl, 0.5% Triton-X-100, 1X cOmpleteTM EDTA-free protease inhibitor (Sigma Aldrich) per 1×106 cells. Cells were agitated in lysis buffer on ice for 30 mins before centrifuging at 12,000RPM, 4°C for 20 mins and removing the supernatant. PierceTM BCA protein assay kit was then used (according to manufacturer’s microplate protocol) to calculate protein concentrations.

For SDS-PAGE gel electrophoresis, required volume of 4x Laemmli buffer (with 10% BME) was added to protein samples and incubated at 95oC for 5 mins. 50µg of protein was then loaded into BIORAD Mini-PROTEAN pre-cast gels (12% acrylamide) and run at 60V for 2hrs. Semi-dry transfer onto nitrocellulose or PVDF membrane was performed (Thermo Scientific Pierce G2 fast blotter) for 15 mins at 15V, 1.3A. Membranes were then blocked in 5% non-fat milk for 1hr at RT before adding primary antibodies to desired concentrations and incubating on a roller overnight at 4oC. The following day, membranes were washed 3x with TBS-T before incubating in secondary antibodies in TBS-T for 1hr at RT. Membranes were then washed 2x with TBS-T and 3x with TBS before imaging on LiCor Odyssey CLx and processing on ImageStudio software.

### RT-qPCR

RNA for qPCR was collected from cell pellets using QIAGEN RNeasy kit according to manufacturer’s protocol. RT-qPCR was then performed using SensiFAST™ SYBR® Lo-ROX One-Step Kit with half volumes according to manufacturer’s protocol and run using Applied Biosystems QuantStudio6 Flex. Primer sequences used were as follows:

*Flotillin-2 (*Forward 5’-GGCTTGTGAGCAGTTTCTGG-3’); (Reverse 5’-AATCTGCTCCACTGTCAGGG-3’), *GAPDH* (Forward 5’-GTCTCCTCTGACTTCAACAGCG-3’); (Reverse 5’-ACCACCCTGTTGCTGTAGCCAA - 3’).

### Antibody Staining and Image Acquisition

Cells were plated onto glass coverslips and following indicated treatment/incubation, cells were washed in 1xPBS and fixed using 4% PFA (10 mins, RT) or modified MEM-Fix (4% formaldehyde, 0.25-0.5% glutaraldehyde, 0.1M Sorenson’s phosphate buffer, pH7.4) (Bodeen *et al*., 2017; Rogers and Scholpp, 2021) for 7 mins at 4°C. Cells were then incubated in permeabilisation solution (0.1% Triton-X-100, 5% serum, 0.1M glycine in 1xPBS) for 1hr at RT. Primary antibodies were diluted in incubation buffer (0.1% Tween20, 5% serum in 1xPBS) and coverslips incubated in 50 µl spots on parafilm overnight at 4°C. Coverslips were then washed with 1xPBS 3x for 5 mins before incubation in 50 µl spots of secondary antibodies diluted in incubation buffer for 1hr at RT. Coverslips were then washed 3x for 5 mins with 1xPBS before mounting onto glass slides using ProLong Diamond anti-fade mountant (Invitrogen) and left to dry for 24hrs before imaging. Confocal microscopy for immunofluorescent antibody imaging was performed on an inverted Leica TCS SP8 X laser-scanning microscope using the 63x water objectives.

### BrdU Proliferation Assay

AGS and HFE-145 cells were co-cultivated on glass cover slips in 6 well plates. After 45 hrs co-cultivation, cells were incubated in media containing 10µg/ml BrdU (Abcam, ab142567) for 3 hrs. Cells were then fixed with 4% PFA for 15 mins at RT. After washing 2x with PBS, immunostaining of BrdU was performed using a BrdU immunohistochemistry kit (Abcam, ab125306) according to manufacturer’s protocol. Cover slips were then mounted onto slides using Prolong Diamond Antifade mountant and left to dry for 24 hrs. Slides were then imaged on a light microscope with 20x objective and captured using an Olympus EP50 colour camera. BrdU-stained cells (brown) were counted and quantified as a percentage of the total cell population (counterstained blue).

### Zebrafish microinjections and image analysis

For experiments in zebrafish embryos, indicated DNA plasmids were injected at the 1-2 cell stage at 100 ng/µl after dechorination. Embryos were left to develop at 28°C until 8 hpf (shield stage). For confocal microscopy analysis, live zebrafish embryos were embedded in 0.7% low melting agarose (Sigma-Aldrich) dissolved in 1x Ringer’s solution. Images of embryos were obtained with an upright Leica TCS SP8X microscope equipped with hybrid detectors (HyD) using 63x dip-in objective.

### In situ hybridisation (ISH)

Pax6a digoxigenin (DIG) antisense RNA probes were generated from linearised plasmids using an RNA labelling and detection kit (Roche) (Scholpp and Brand, 2003). Embryos for ISH were dechorinated and injected with indicated DNA plasmids (100ng/µl) at the 1-2 cell stage. Embryos were left to develop at 28oC for 30 hrs before fixation in 4% PFA overnight at 4°C. Embryos were then washed twice in PBST and dehydrated in 100% methanol for 30 mins at RT. Following 2x PBST wash, they were re-fixed in 4% PFA for 30 mins and washed again 2x. Embryos were then incubated in Hyb+ solution at 69°C for 4-6 hrs before replacing the solution with probe-Hyb+ mix (1:20 Pax6a probe in Hyb+) and incubated overnight. The next day, embryos underwent several washing steps in 25% Hyb-, Hyb-, 2xSSCT and 0.2xSSCT solutions at 69°C, then MABT at RT. Embryos were then incubated in 2% blocking buffer for 4-5 hrs, followed by 1:4000 anti-DIG antibody (Roche) in 2% blocking buffer overnight at 4°C. The next day, embryos were washed 5x 15 mins in MABT and then once in NTMT (5 mins). Embryos were transferred to 24 well plates and NTMT-NCP-BCIP mix (1:200 dilution NCP-BCIP:NTMT (Roche)) was added. Embryos were left to develop signal staining for 2 hrs in the dark (RT) prior to NTMT and PBST washes and 30 min re-fix in 4% PFA (RT). Two final washes in PBST were performed before storing embryos in 70% glycerol. Stained embryos were then imaged on a stereo microscope and measurements of the forebrain and midbrain made using Fiji software.

### Quantifications and Statistical Analyses

All filopodia quantifications were calculated from Z-stack images of cells expressing membrane-mCherry and were done using Fiji software. Filopodia length was measured from the tip of the filopodia to the base, where it contacted the main cell body. In the case of branching protrusions, one branch (the longest) would be measured. Quantifications of Western blot images was performed by measuring the mean gray value of bands (after subtracting background) and normalising to loading controls. Quantification of qPCR data used the Pfaffl equation to calculate fold-change expression from Ct values, after normalising to GAPDH. All experiments / conditions were repeated at least in biological triplicates. Significance was tested using Mann-Whitney U or Kruskal-Wallis Tests (non-parametric) or student’s t-test (parametric) against the relative controls. Bonferroni correction for multiple comparisons was applied. Error bars on bar charts show standard error of the mean (SEM).

## Supplementary Figure legends

**Supplementary Figure 1.**
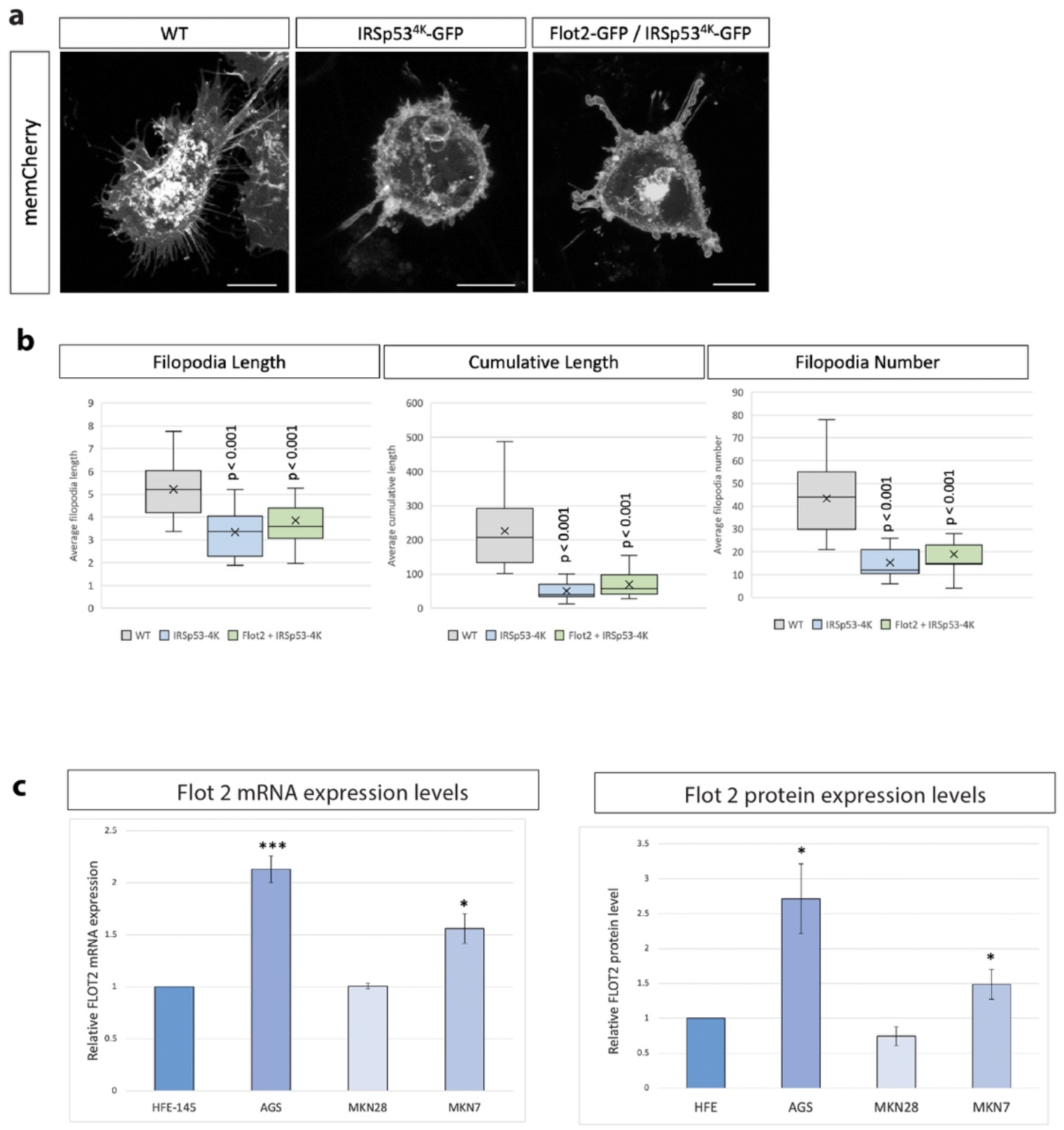
**a,** Confocal images of AGS cells expressing the dominant negative IRSp53^4K^-GFP mutant and memCherry, showing inhibition of filopodia formation (even in the presence of Flot2-GFP). Scale bar 10µm. **b,** Quantification of filopodia of cells from (a). (n per condition = 25, 13, 13; n = number of cells). Significance calculated by Student’s t-test. **c,** Quantification of Flot2 expression levels (mRNA and protein) in HFE-145, AGS, MKN7, and MKN24 cells. Significance calculated by Student’s t-test.

**Supplementary Figure 2.**
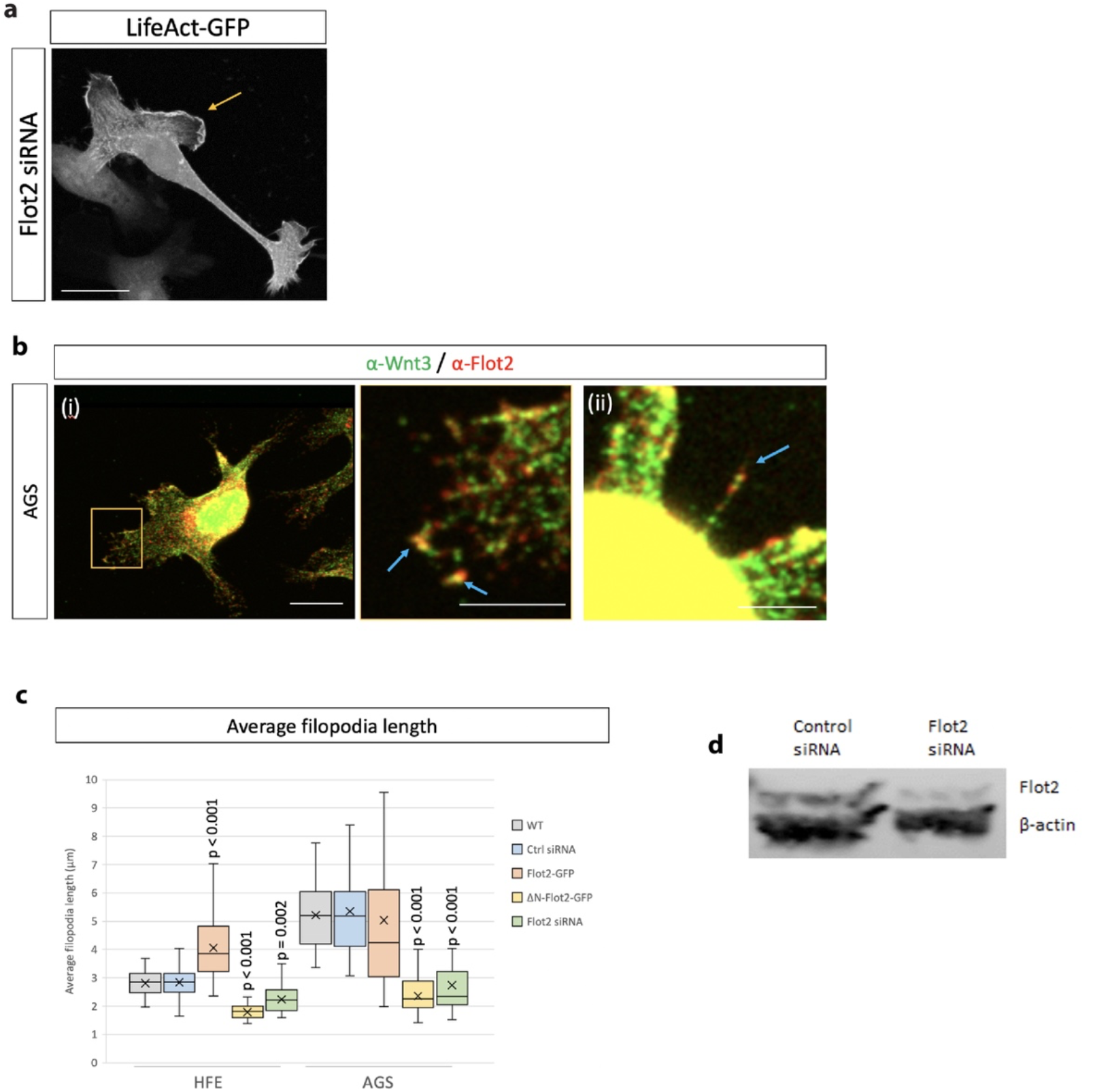
Flotillin-2 regulates filopodia and lamellipodia formation in AGS cells. **a**, Confocal image of an AGS cell with depleted Flot2 (siRNA-mediated) and expressing LifeAct-GFP to visualise actin. Yellow arrow highlights example of lamellipodia frequently seen in Flot2-depleted cells. Scale bar 10µm. **b,** Immunofluorescent images of AGS cells stained for Wnt3 (green) and Flot2 (red). Co-localisation of Flot2 and Wnt3 highlighted by blue arrows. Scale bars represent (i) 10µm (left) and 5µm (right), (ii) 5µm. **c,** Quantification of filopodia length in HFE and AGS cells expressing indicated constructs. Significance calculated by student’s t-test with Bonferroni correction for multiple comparisons. **d,** Analysis of efficiency of siRNA-mediated knock-down of Flot2 in AGS cells.

**Supplementary Figure 3.**
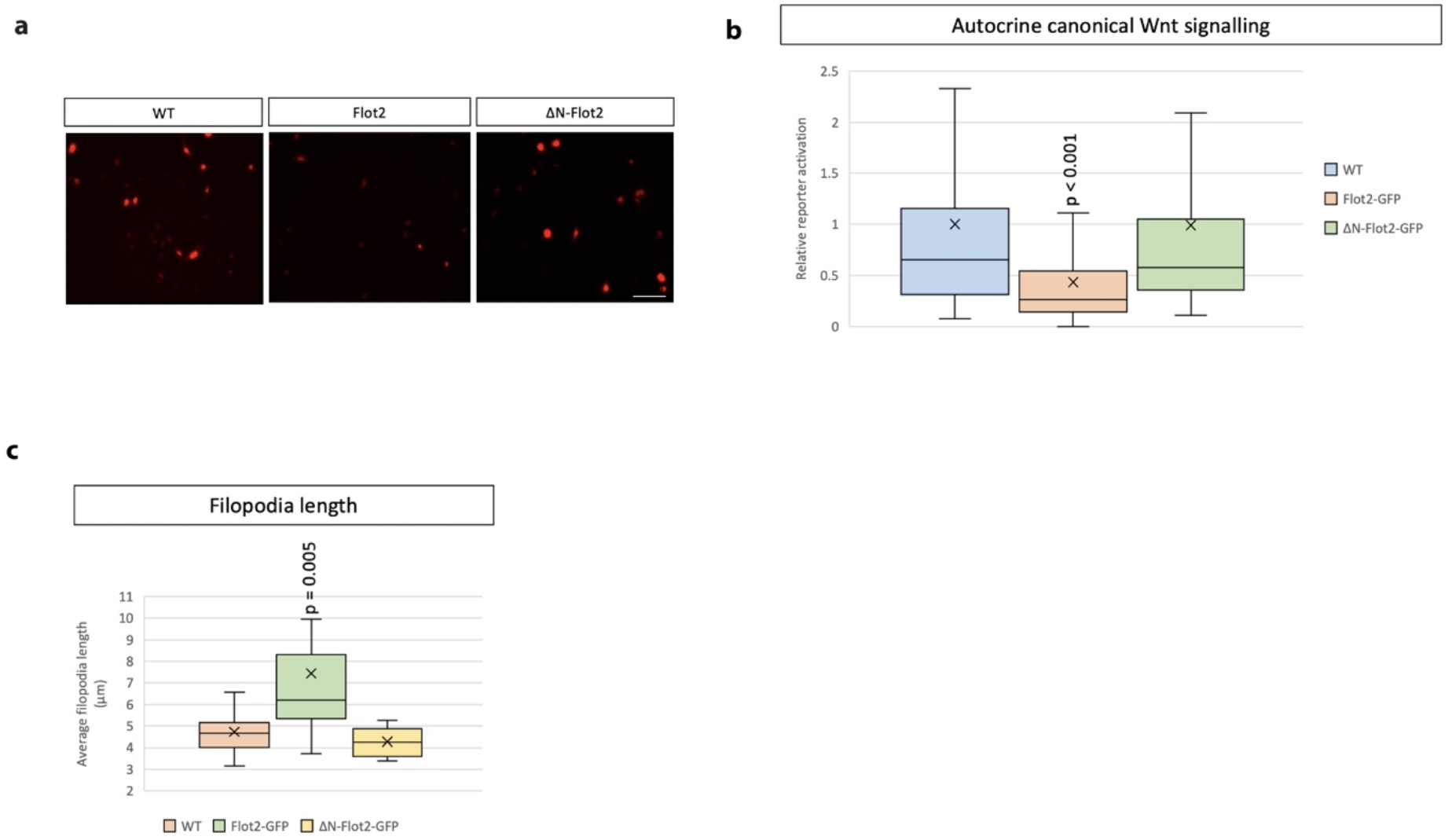
Flotillin-2 inhibits autocrine canonical Wnt signalling. **a,** Representative images of AGS cells transfected with the 7xTCF-NLS-mCherry reporter and indicated constructs for 48 hrs. Scale bar 20µm. **b,** Relative quantification of 7xTCF-NLS-mCherry fluorescence compared to untransfected control. Significance calculated by student’s t-test. (n per condition = 406, 212, 469; n = number of nuclei measured). **c,** Quantification of filopodia length in zebrafish epiblast cells at 8 hpf after microinjection of indicated constructs (at 100ng/µl). Significance calculated by student’s t-test.

